# Lithium treatment reverses irradiation-induced changes in rodent neural progenitors

**DOI:** 10.1101/579235

**Authors:** Zanni Giulia, Goto Shinobu, Gaudenzi Giulia, Naidoo Vinogran, Levy Gabriel, Di Martino Elena, Dethlefsen Olga, Cedazo-Minguez Angel, Merino-Serrais Paula, Hermanson Ola, Blomgren Klas

## Abstract

Cranial radiotherapy in children has detrimental effects on cognition, mood, and social competence in young cancer survivors. Treatments harnessing hippocampal neurogenesis are currently of great relevance in this context, and we previously showed that voluntary running introduced long after irradiation rescued hippocampal neurogenesis in young mice (Naylor et al. 2008a). Lithium, a well-known mood stabilizer, has both neuroprotective, pro-neurogenic as well as anti-tumor effects, and in the current study we introduced lithium treatment 4 weeks after irradiation, analogous to the voluntary running study. Female mice received a single 4 Gy whole-brain irradiation dose at postnatal day (PND) 21 and were randomized to 0.24% Li_2_CO_3_ chow or normal chow from PND 49 to 77. Hippocampal neurogenesis was assessed at PND 77, 91 and 105. We found that lithium treatment had a pro-proliferative effect on neural progenitors and promoted neuronal integration upon its discontinuation. Gene expression profiling and DNA methylation analysis identified two novel factors related to the observed effects, Tppp, associated with proliferation, and GAD2/65, associated with neuronal signaling. Our results show that lithium treatment reverses irradiation-induced impairment of hippocampal neurogenesis even when introduced long after the injury. We propose that lithium treatment should be intermittent in order to first make neural progenitors proliferate and then, upon discontinuation, allow them to differentiate. Our findings suggest that pharmacological treatment of cognitive so-called late effects in childhood cancer survivors is possible.

## INTRODUCTION

Dramatic improvements in childhood cancer survival rates have been made in the last decades (Gatta et al. 2014) and this is due to great strides in the intervention treatments. The treatments encompass a combination of surgery, chemotherapy and radiotherapy, with the recent addition of immunotherapy. However, the growing population of survivors often has to face the therapy-related morbidity (Spiegler et al. 2004). Radiotherapy is known to cause debilitating cognitive alterations (Georg Kuhn and Blomgren 2011) leading to impaired processing speed, attention and working memory, further impinging on emotional and psychological well-being and ultimately leading to anxiety, and posttraumatic stress like symptoms (PTSS) (Marusak et al. 2017). Declines in IQ and academic achievements have been observed during longitudinal follow-up and this impact the quality of life, school attendance and overall daily activities (Janelsins et al. 2011).

Different mechanisms have been implicated in the cognitive changes observed in patients treated with radiotherapy (Davis et al. 2013). Increased levels of cytokines seem to mediate some of these changes, as well as direct and indirect DNA damage, endocrine dysfunction, activation of microglia and astrocytes, hypomyelination and decreased neurogenesis (Janelsins et al. 2011) (di Fagagna et al. 2003) (Monje et al. 2002). In particular neurogenic regions, harboring cellular proliferation, display higher sensitivity to irradiation as seen in rodent models and in humans (Fukuda et al. 2004) (Limoli et al. 2004). Irradiation, even after a single moderate dose, was shown to cause apoptosis and progressive decline in neurogenesis of young rats and mice resulting in severe cognitive declines (Fukuda et al. 2004) (Boström et al. 2013). Adult hippocampal neurogenesis persists throughout life, mainly in the subgranular zone (SGZ) of the hippocampus and in the subventricular zone (SVZ) of the lateral ventricles (Altman and Das 1965). These regions harbor neural stem and progenitor cells (NSPCs) dividing continuously and giving birth to newborn neurons. This is believed to contribute highly to hippocampal plasticity and especially learning, memory and mood regulation (Shors et al. 2002) (Deng, Aimone, and Gage 2010) (Yun et al. 2016).

Lithium, commonly used in the treatment of bipolar disorder, has been shown to exert neuroprotective and regenerative effects in a variety of neurological insults (Shorter 2009). In preclinical studies lithium protected the neonatal brain against the neurodegenerative effects of hypoxia-ischemia (HI) (Xie et al. 2014) and rescued cognitive loss in adult as well as in young mice after cranial irradiation (Yazlovitskaya et al. 2006) (Huo et al. 2012; Zhou et al. 2017). The neuroprotective effects of lithium after cranial radiation are attributable to enhanced hippocampal neurogenesis and decreased apoptosis in young rats and mice (Yazlovitskaya et al. 2006) (Huo et al. 2012). Lithium also restored synaptic plasticity in a Down syndrome mouse model (Contestabile et al. 2013) and ongoing trials aim at introducing lithium as a treatment of a broad range of brain related disorders (“Neuroprotective Effects of Lithium in Patients With Small Cell Lung Cancer Undergoing Radiation Therapy to the Brain - Full Text View - ClinicalTrials.gov” n.d.).

Despite the surge of studies conducted on lithium, the exact mechanisms of action are still only partly elucidated. Lithium exerts its action through modulation of intracellular second messengers with subsequent alteration of complex and interconnected intracellular enzyme cascades (Brown and Tracy 2013). One target is the protein kinase Gsk3β (O’Brien and Klein 2009). Direct (Wang et al. 2015) (Jope 2003) and indirect (Zhang et al. 2003) inhibition of Gsk3β by lithium and related improvements of impaired cognition likely involve a variety of different mechanisms, such as supporting long-term potentiation and diminishing long-term depression, promotion of neurogenesis and ultimately reduction of inflammation and apoptosis (Yazlovitskaya et al. 2006) (Huo et al. 2012) (King et al. 2014) (Voytovych, Kriváneková, and Ziemann 2012).

Encouraging results also support the use of lithium in combination with cancer treatment to improve the therapeutic effect, for example as a radiosensitizer (Zhukova et al. 2014) (Ronchi et al. 2010) (Zinke et al. 2015) (Korur et al. 2009). Nevertheless, a post radiotherapy lithium treatment may still be preferable to safely exclude the likelihood of protecting tumor cells and thus increasing the risk of relapses. The study herein investigates the effects of lithium treatment on NSPC proliferation, dendritic orientation and survival after brain irradiation and whether using a therapeutically relevant dose can rescue neurogenesis even long after irradiation of the juvenile brain. Ultimately, we provide evidence for a possible molecular mechanism involving novel proteins targeted by lithium and further elucidate the effects of irradiation on fate commitment of NSPCs and how lithium can harness this process.

## RESULTS

Assessment of neurogenesis (proliferation, dendritic orientation and survival of the NSPCs) was performed at three different time-points, immediately after (PND 77), two weeks after (PND 91) and 4 weeks after (PND 105) termination of LiCl exposure (**Figure 1A** *in vivo* study design).

**Figure 1.**
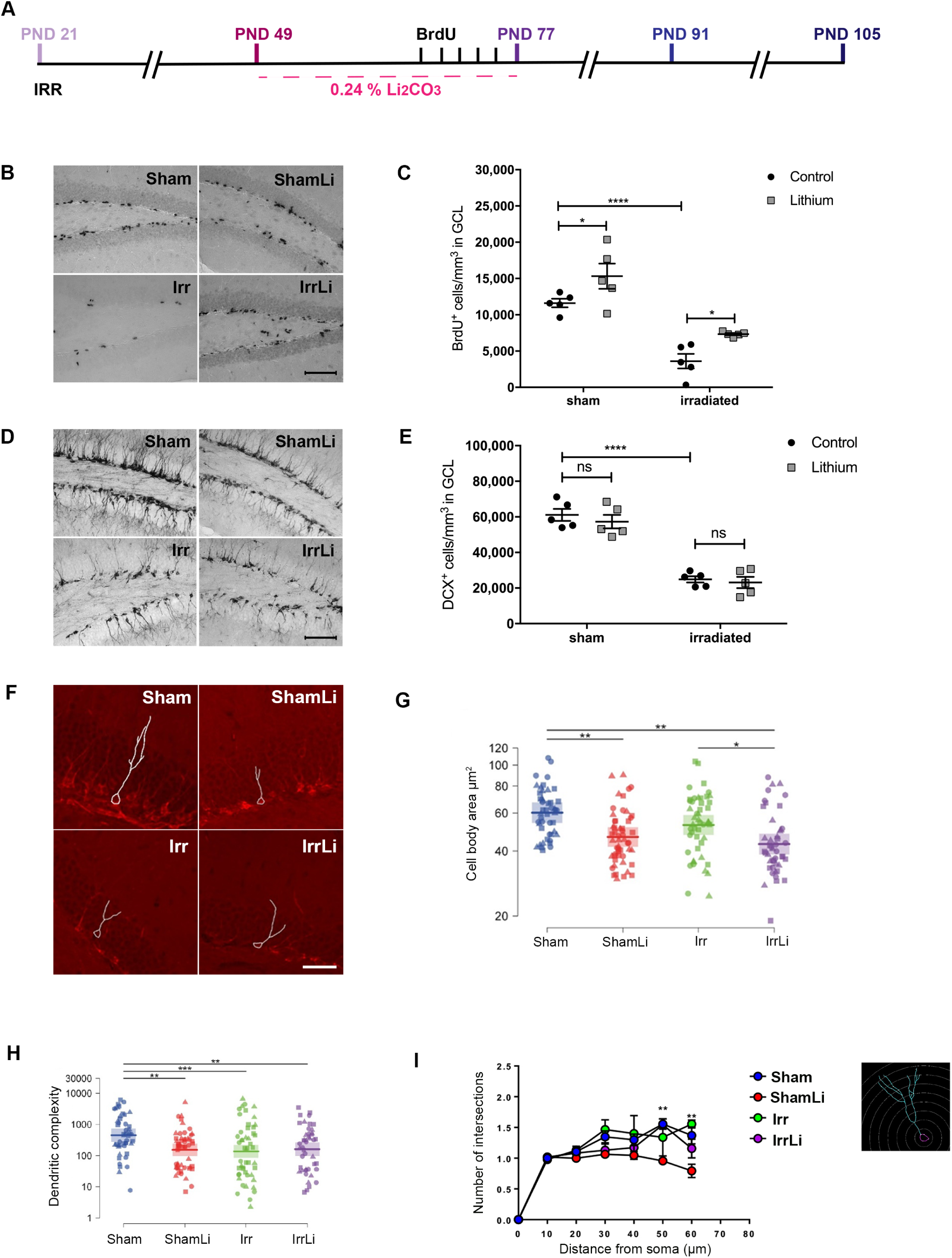
Delayed lithium treatment after irradiation increases proliferation to a halt of neuronal maturation in the GCL at PND77. **(A)** Timeline of the *in vivo* experimental design showing that lithium treatment was delayed in respect to irradiation and 3 different time-points (PND 77, 91 and 105) were investigated to assess the early as well as the late effects of lithium on neurogenesis. **(B)** BrdU immunoreactivity in the GCL of the different treatment groups at PND77. Scale bar= 100 µm. 20x magnification. **(C)** Interleaved dot plot graph of BrdU quantification in GCL showing the effect of lithium treatment in the sham (\**p*=0.0485) and irradiated group (\**p*=0.0471). 2-way ANOVA shows that both irradiation and lithium treatments have an effect: irradiation (F_1,16_=57.77, \*\*\*\**p*<0.0001), LiCl (F_1,16_=12.45, \*\**p*=0.0028). **(D)** DCX immunoreactivity in the GCL of the different treatment groups at PND77. Scale bar=100 µm. 20x magnification. **(E) I**nterleaved dot plot graph of DCX quantification in the GCL at PND77. 2-way ANOVA resulted in an effect of irradiation but not lithium treatment: irradiation (F_1,16_=127.3, *\*\*\*\*p*<0.0001), LiCl (F_1,16_=1.34, *p*=0.7974). Lithium treatment did not alter the density of DCX positive cells in sham (*p*=0.8097) and irradiated (*p*>0.9999) groups. **(F)** Representative images of the dendritic tracing in DCX+ cells for all experimental groups. **(G)** Cell body area (µm^2^) shows that lithium treatment significantly decreased the size of the immature cells both in sham and irradiated groups at PND77. **(H)** Dot plot graph of the dendritic complexity of DCX+ immature neurons shows significant decrease following lithium treatment, irradiation, and lithium treatment after irradiation compared to sham at PND77. Circle, triangle and square marks represent the value for each neuron in each animal. N (number of animals) = 3 in each group. Number of traced neurons is 15 to 20 in each animal. Linear mixed model, * *p* < 0.05, ** *p* < 0.01, *** *p* < 0.001. Error bars represent point estimate and 95% confidence interval. **(I)** Sholl analysis of DCX+ immature neurons shows that lithium treatment reduces the number of intersections in the distal part (50-60 µm) of the dendrites at PND77 in sham but not in irradiated treated groups. Error bars represent SEM ****p<0.0001; **p<0.01; *p<0.05.

### Four Weeks of Continuous Lithium Treatment Increases the Number of Proliferating Cells in the DG but Decreases the Cell Body Area and the Dendritic Complexity of DCX^+^ Cells

To analyze the effect of lithium on proliferation of the NSPCs, we measured BrdU at PND 77 (**Figure 1B**). Proliferation in the GCL was decreased significantly by irradiation in both the control and the lithium-treated mice (**Figure 1C**). Lithium increased the density of proliferating cells in both sham and irradiated brains, around 83% in the irradiated group and 33% in the sham group. We further determined the effects of lithium on the density of doublecortin-positive (DCX^+^) cells in the GCL at PND 77 (**Figure 1D**). Irradiation provoked a 60% decrease of the density of DCX^+^ cells in the GCL, but lithium had no effect on DCX^+^ cell density, neither in sham, nor in irradiated brains (**Figure 1E**).

In addition, we conducted morphometric analysis of DCX^+^ immature neurons at PND 77 (**Figure 1F**). We found that lithium treatment decreased the cell body area in both sham and irradiated animals (**Figure 1G**), whereas radiation drastically decreased the dendritic complexity. Surprisingly, lithium treatment *per se* reduced dendritic complexity in sham, but not in irradiated animals at PND 77 (**Figure 1H**). This was confirmed by Sholl analysis, showing that lithium treatment reduced the number of intersections at the distal part of the dendrites (50 and 60 mm from the soma) at PND 77 in the sham, but not in irradiated animals (**Figure 1I**).

### Lithium Discontinuation Prevented Irradiation-Induced Alterations in Process Orientation and Dendritic Complexity of DCX^+^ Cells at PND 91

To assess if the increase in proliferating cells at PND 77 resulted in increased survival and differentiation into immature neurons, we analyzed the density of DCX^+^ cells in the GCL at PND 91, two weeks after lithium discontinuation (**Figure 2A**). DCX^+^ cell density was decreased by irradiation, as expected, but this decrease was reversed by lithium treatment. Also in non-irradiated brains the DCX^+^ cell density increased. The increase was 156% and 24% in the irradiated and sham lithium treated groups, respectively (**Figure 2B**). To further address the effects of radiation and lithium treatment on the integration of newly born neurons, we performed a DCX/BrdU double immunostaining (**Figure 2C**), enabling us to determine if the orientation of the main dendritic process of neurons born during the last 5 days of lithium treatment was parallel or radial to the GCL. We have earlier shown that irradiation causes the main dendritic process to shift from a radial to a parallel orientation (Naylor et al., 2008, PNAS). Our results confirmed that irradiation increased the percentage of parallel processes from 22% in sham to 41% in irradiated, whereas the number of radial processes was decreased from 64% in sham to 48% in irradiated (**Figure 2D**). Remarkably, we found that lithium treatment reduced the percentage of parallel main processes by half, from 41% to 21%, in the irradiated brains, to the same level as in the sham controls (22%) (**Figure 2D**). However, lithium did not alter the proportion of radial main processes, neither in the sham, nor in the irradiated brains. Also, we found that lithium treatment increased the number of BrdU cells that were not labelled for DCX in the irradiated group (25%) compared to the sham irradiated group (11%). However, the morphometric analysis of the DCX^+^ cells at PND 91 (**Figure 2E**) revealed that the cell body area of the irradiated groups were not different from the sham groups, and lithium did not affect cell body area either (**Figure 2F**). The dendritic complexity of the DCX+ cells in the irradiated group was drastically decreased compared to the sham group (7%), but lithium treatment restored this completely to the sham level (**Figure 2G**). This was further confirmed by Sholl analysis, where we found that the number of intersections at 50 and 60 µm from the soma was normalized to sham level in the irradiated group treated with lithium at PND 91 (**Figure 2H**).

**Figure 2.**
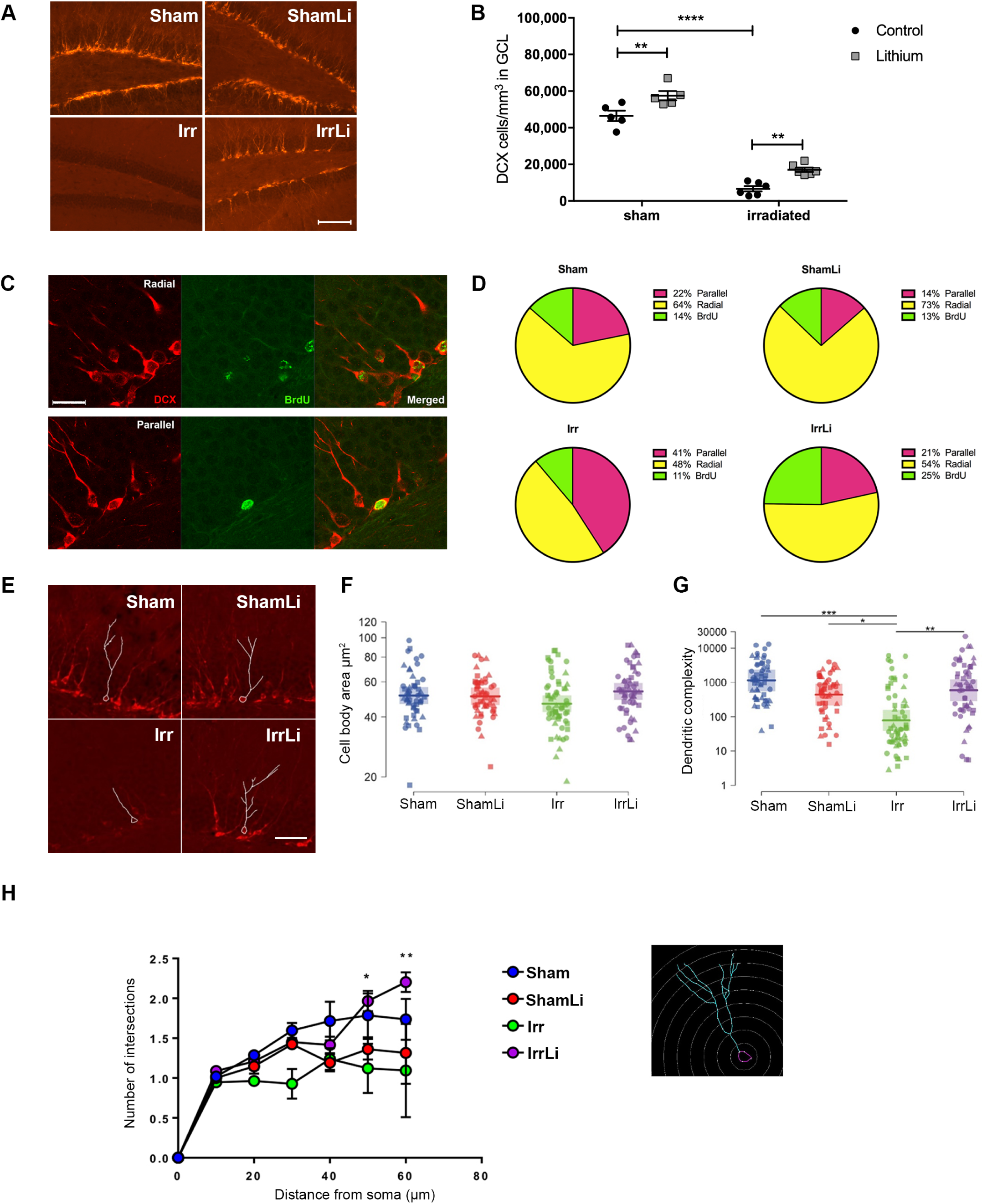
Lithium restores irradiation-induced alteration in dendrite arborization of doublecortin (DCX) cells in the subgranular zone (SGZ) of the DG at PND91. **(A)** DCX immunoreactivity in the GCL of the different treatment groups at PND91. Scale bar=100 µm. 20x magnification. **(B)** Interleaved bar graph of the quantification of DCX^+^ cells in the GCL showing the effect of lithium treatment in the sham group (\*\**p*=0.0030) and in the irradiated group (\*\**p*=0.0023). 2-way ANOVA shows the effect of both treatments: irradiation (F_1,18_=400.6, \*\*\*\**p*<0.0001), LiCl (F_1,18_=28.70, \*\*\*\**p*<0.0001). **(C)** The phenotype of the radial (upper panel) and parallel (lower panel) processes in immature neurons at PND91 was determined by double labeling for BrdU and DCX as shown in the representative confocal images of DCX+ (red), BrdU+ (green) and merged BrdU+-DCX+ cells in the GCL. Scale bar=20 µm. 63x magnification. **(D)** Pie chart of the percentages of BrdU cells double labeled for DCX and with either radial or parallel process phenotype. The percentages of radial processes were not affected by lithium treatment: in the sham group *p*=0.2341 and in the irradiated group *p*=0.5313. 2-way ANOVA shows the effect of irradiation on radial processes: irradiation (F_1,18_=24.4, \*\*\**p=*0.0001), LiCl (F_1,18_=3.96, *p=*0.0619). The percentages of parallel process were increased after irradiation and lithium treatment restored the percentages of parallel processes in the irradiated group (\*\*\*\**p*<0.0001) but not did not affect the sham group (*p*=0.0570). 2-way ANOVA: irradiation (F_1,18_=33.59, \*\*\*\**p*<0.0001), LiCl (F_1,18_=34.75, \*\*\*\**p*<0.0001). The percentages of BrdU not labeled for DCX were not affected by irradiation but were increased in the lithium treated irradiated group (\*\**p*=0.0094). 2-way ANOVA: irradiation (F_1,18_=2.29, *p*=0.1476), LiCl (F_1,18_=4.212, *p*=0.0550). **(E)** Representative images of the dendritic tracing in DCX+ cells in animals for all experimental groups. **(F)** Dot plot graph of the cell body area showing no effect of either lithium or irradiation at PND91. **(G)** Dot plot graph of the dendritic complexity of DCX+ immature neurons shows significant decrease in irradiated treated group compared to all the other groups at PND91. Circle, triangle and square marks represent the value for each neuron in each animal. N (number of animals) = 3 in each group. Number of traced neurons is 15 to 20 in each animal. Linear mixed model, * *p* < 0.05, ** *p* < 0.01, *** *p* < 0.001. Error bars represent point estimate and 95% confidence interval. **(H)** Sholl analysis of DCX+ immature neurons shows that lithium treatment restores the number of intersections in the distal part (50-60 µm) of the dendrites at PND91 in the irradiated but not in the sham treated group. N (number of animals) = 3 in each group. Number of traced neurons is 15 to 20 in each animal. Two-way ANOVA and post-hoc Bonferroni test. \**p*<0.05, \*\*\**p*<0.001 compared to irradiation only group. Error bars represent SEM. ****p<0.0001; **p<0.01; *p<0.05.

### Lithium Upregulates Genes Involved in Cell Cycle and Signaling after Irradiation *in Vitro* – association with changes in DNA methylation

In order to investigate the molecular mechanisms possibly involved in the lithium effects after irradiation, we designed an *in vitro* study (**Figure 3A**), where isolated NSPCs from the brains of 15.5 days rat embryos, after expansion passage 3 (P3) were exposed to 2.5 Gy irradiation and lithium treatment. Initially, we performed RNA sequencing and compared the reactome across different conditions. Principal component analysis (PCA) revealed that the sham groups clustered together for PC2 (**Figure 3B**) whereas the irradiated groups clustered for PC3 (**Figure 3C**), indicating that NSPCs responded in a specific manner to the individual treatments. A deeper analysis of the reactome profile with respect to each PC revealed that PC2 identified changes in gene expression related to a variety of brain developmental as well as synaptic transmission processes (**Figure 3D**), whereas PC3 identified differential gene expression for genes involved primarily in cell cycle as well as axonal guidance (**Figure 3E**). To identify individual candidate genes involved in the specific responses, we next looked for the highest fold-changes in gene expression between sham and sham treated with lithium (**Figure 3F**) and between irradiated and irradiated treated with lithium (**Figure 3G**). This analysis highlighted two genes, one that encodes for tubulin polymerization-promoting protein (Tppp) and the glutamate decarboxylase 2 (*GAD2*) gene that encodes for the GAD65 protein (Erlander et al. 1991). The mRNA levels were confirmed by RT-qPCR and lithium accounted for most of the variability in the sham and irradiated groups in Tppp expression levels (**Figure 3H**) as well as in GAD65 (**Figure 3I**).

**Figure 3.**
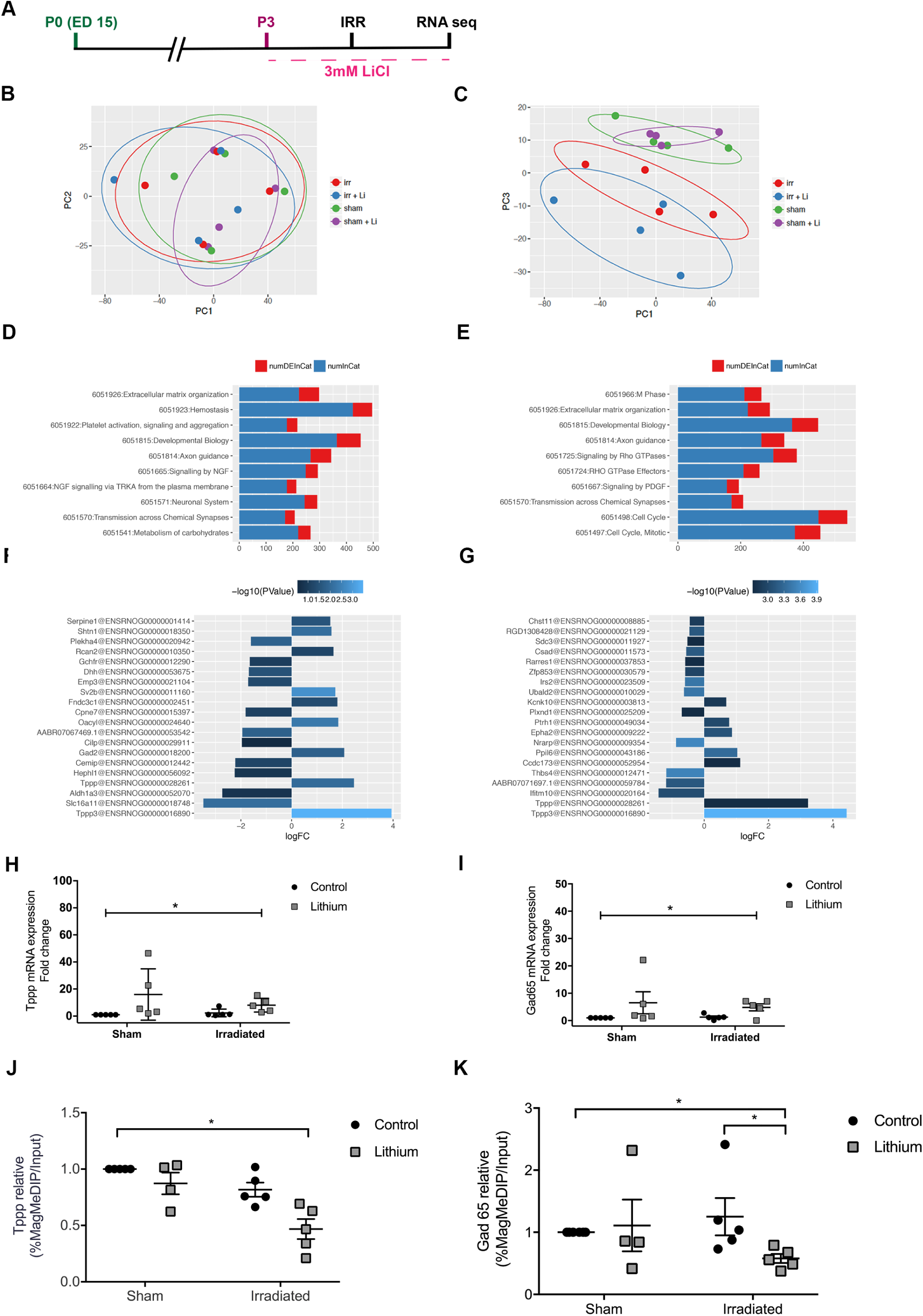
Lithium regulates the expression of genes involved in cell cycle and neuronal transmission *in vitro*. **(A)** Timeline of the *in vitro* experimental design showing that neural stem progenitor cells (NSPC) at passage 3 (P3) were exposed to lithium and irradiation followed by RNA sequencing. **(B)** and **(C)** Principal component (PC) analysis plot based on the filtered and normalized counts per million that allowed us to identify the uncorrelated variables to explain the source of maximum amount of variance in the different treatment groups (sham, shamLi, irradiated and irradiated lithium). **(D)** Gene ontology (GO) analysis revealed that the changes in the reactome of different treatment groups for PC2 were related to neuronal transmission. ‘numDEInCat’ is the number of differentially expressed (DE) genes in the corresponding GO category, whereas ‘numInCat’ is the number of detected genes in the corresponding GO category. **(E)** Gene ontology (GO) analysis revealed that the changes in the reactome of different treatment groups for PC3 were related to cell cycle regulation. **(F)** Table of the differentially expressed genes in Sham compared to Sham treated with lithium groups. **(G)** Table of the differentially expressed genes in Irradiated compared to Irradiated treated with lithium groups. **(H)** Gene expression analysis of the identified cell cycle gene shows that lithium significantly increases Tppp expression in both sham and irradiated groups. 2-way ANOVA shows the effect of lithium treatment: irradiation (F_1,16_=0.5473, *p*=0.4702), LiCl (F_1,16_=5.439, \**p*=0.0331). **(I)** Gene expression analysis of the identified neuronal transmission gene shows that lithium significantly increases GAD65 expression in both sham and irradiated groups. 2-way ANOVA shows the effect of lithium treatment: irradiation (F_1,16_=0.1073, *p*=0.7475), LiCl (F_1,16_=4.576, \**p*=0.0482). Error bars represent SEM. ****p<0.0001; **p<0.01; *p<0.05. **(J)** Methylation analysis by MeDIP-qPCR of Tppp showing the significant difference between groups (\**p=*0.0145). Post hoc test shows the significant decrease of Irr+li group compared to the Sham group (\**p=*0.0123). **(K)** MeDIP-qPCR analysis of Gad65 showing the significant difference between groups (\**p=*0.0112). Post hoc test shows the significant decrease of Irr+li group compared with Sham (\**p=*0.0284) and Irr+li group (\**p=*0.0273).

To investigate possible mechanisms underlying the changes in gene expression, we next investigated the effects on DNA methylation of the regulatory regions of Tppp and the GAD2 gene, the latter encoding GAD65, using a MeDIP-based approach (Brebi-Mieville et al. 2012) and analyzing the 5-methylcytosine (5mc) levels of the regulatory regions of these two genes in fetal neural stem cells *in vitro*. DNA methylation of regulatory regions is associated with transcriptional repression, and decreased levels of methylation are thus mostly associated with increased gene expression. These experiments revealed a clear correlation between the effects of irradiation and lithium treatment on gene expression and the corresponding DNA methylation levels. At the *Tppp* gene, 5mc levels were significantly decreased after lithium treatment of irradiated cells compared to the sham control (**Figure 3J**). The 5mc levels at the *GAD2* regulatory region also showed a significant decrease in the irradiated group treated with lithium compared to sham control and irradiated groups (**Figure 3K**). These results suggest that lithium treatment after irradiation influences epigenetic mechanisms yielding decreased DNA methylation levels.

### Lithium Upregulates Tppp and GAD65 levels in the Irradiated Juvenile Brain *in Vivo*

To be able to validate the DNA methylation and RNA sequencing data in our *in vivo* model, we investigated the protein expression levels of Tppp and GAD65 in hippocampal tissue by immunoblotting. The levels of Tppp at PND 77 were the same in all treatment groups (**Figure 4A, B and C**), whereas at PND 91 a significant increase in Tppp expression was observed in the irradiation lithium-treated group (**Figure 4D, E and F**). The levels of GAD65 at PND 77 were significantly reduced in the irradiation group but unaltered in the others (**Figure 4G, H and I**), however at PND 91 its expression increased significantly in the irradiation group treated with lithium (**Figure 4J, K and L**).

**Figure 4.**
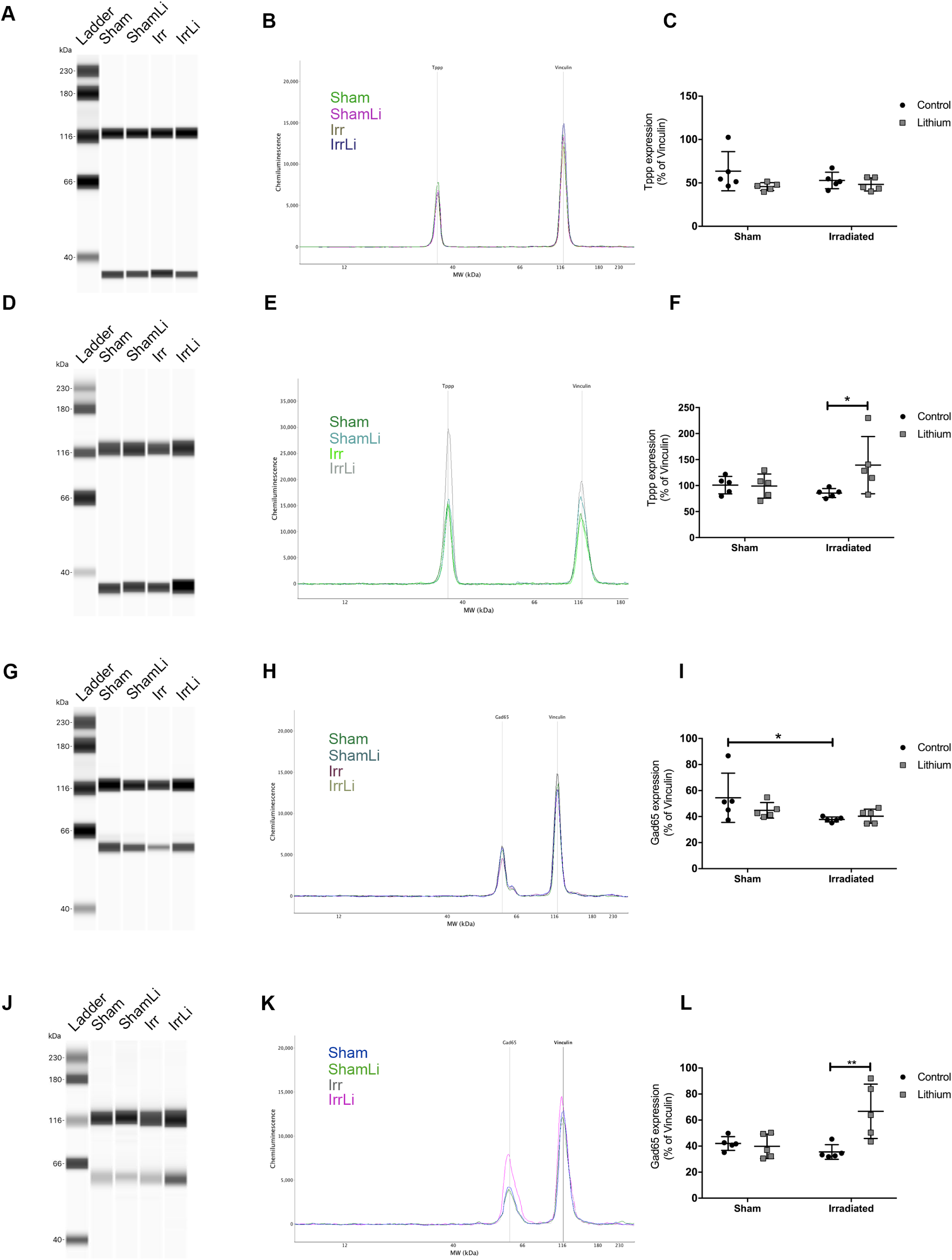
Lithium regulates the expression of cell cycle and neuronal transmission proteins *in vivo* in the mouse DG. **(A)** Representative western blot lane of Tppp migration at PND77. Tppp signal was observed at the MW of 37 kDa. Vinculin used as loading control was observed at the expected MW of 118 kDa. **(B)** Representative chemiluminescence graph showing the peak of both Tppp and Vinculin for each treatment group at PND77. **(C)** Dot plot graph showing the quantification of the normalised chemiluminescence peak of Tppp against its loading control Vinculin at PND77. Tppp expression level in the DG was unaltered in all treatment groups. 2-way ANOVA: irradiation (F_1,16_=0.4998, *p*=0.4898), LiCl (F_1,16_=3.654, *p*=0.0740). **(D)** Representative western blot lane of Tppp migration at PND91. **(E)** Representative chemiluminescence graph showing the peak of both Tppp and Vinculin for each treatment group at PND91. **(F)** Dot plot graph showing the quantification of the normalised chemiluminescence peak of Tppp against its loading control Vinculin at PND91. Tppp expression level in the DG was significantly increased by lithium in the irradiated group (\**p*=0.0313). 2-way ANOVA: irradiation (F_1,16_=0.7927, *p*=0.3865), LiCl (F_1,16_=3.414, *p*=0.0832). **(G)** Representative western blot lane of GAD65 migration at PND77. GAD65 signal was observed at the MW of 65 kDa. Vinculin used as loading control was observed at the expected MW of 118 kDa. **(H)** Representative chemiluminescence graph showing the peak of both GAD65 and Vinculin for each treatment group at PND77. **(I)** Dot plot graph showing the quantification of the normalised chemiluminescence peak of GAD65 against its loading control Vinculin at PND77. GAD65 expression level in the DG was significantly reduced in the irradiated group (\**p*=0.0431) but no change was observed in the other groups. 2-way ANOVA: irradiation (F_1,16_=5.232, \**p*=0.0361), LiCl (F_1,16_=0.599, *p*=0.4502). **(J)** Representative western blot lane of GAD65 migration at PND91. **(K)** Representative chemiluminescence graph showing the peak of both GAD65 and Vinculin for each treatment group at PND91. **(L)** Dot plot graph showing the quantification of the normalised chemiluminescence peak of GAD65 against its loading control Vinculin at PND91. GAD65 expression level in the DG was significantly increased by lithium in the irradiated group (\*\**p*=0.0017). 2-way ANOVA: irradiation (F_1,16_=3.543, *p*=0.0781), LiCl (F_1,16_=7.228, \**p*=0.0161). Error bars represent SEM. ****p<0.0001; **p<0.01; *p<0.05.

### Lithium Prevents the Irradiation-Induced Changes in NSPC Fate Progression Observed 4 Weeks after Its Discontinuation

To assess the survival and the differentiation of the NSPCs at later time in the neurogenic process, we conducted a triple immunostaining of BrdU, NeuN and S100β in the DG at PND 105 (**Figure 5A**). We found that irradiation drastically reduced the proportion of NeuN/BrdU^+^ cells (from 76% to 40%) and increased the proportion of S100β/BrdU^+^ cells (from 10% to 21%) (**Figure 5B**). Lithium after irradiation was able to restore the percentage of neuronal commitment from 40% to 66% approaching sham levels. We did not observe lithium-induced changes in astrocytic commitment in both sham and non-irradiated animals. In addition, irradiation significantly increased the percentage of surviving neurons (BrdU^+^) that were neither of neuronal or astrocytic lineage. To further assess whether lithium had a long-lasting effect on the positive modulation on DCX^+^ cells, we analyzed the density of the DCX^+^ cells in the GCL at PND 105 (**Figure 5C**). The density was still significantly decreased after irradiation, consistent with previous models (Boström et al. 2013) (**Figure 5D**). Additionally, there was no significant difference in the density of DCX^+^ cells between the control and the lithium-treated mice, neither in the irradiated, nor in the sham groups (**Figure 5D**). Together, this indicates that lithium treatment promotes proliferation of NSPCs and that discontinuation of the treatment promotes a wave of neuronal differentiation, but at the same time prevents the wave of irradiation-induced astrocytic differentiation.

**Figure 5.**
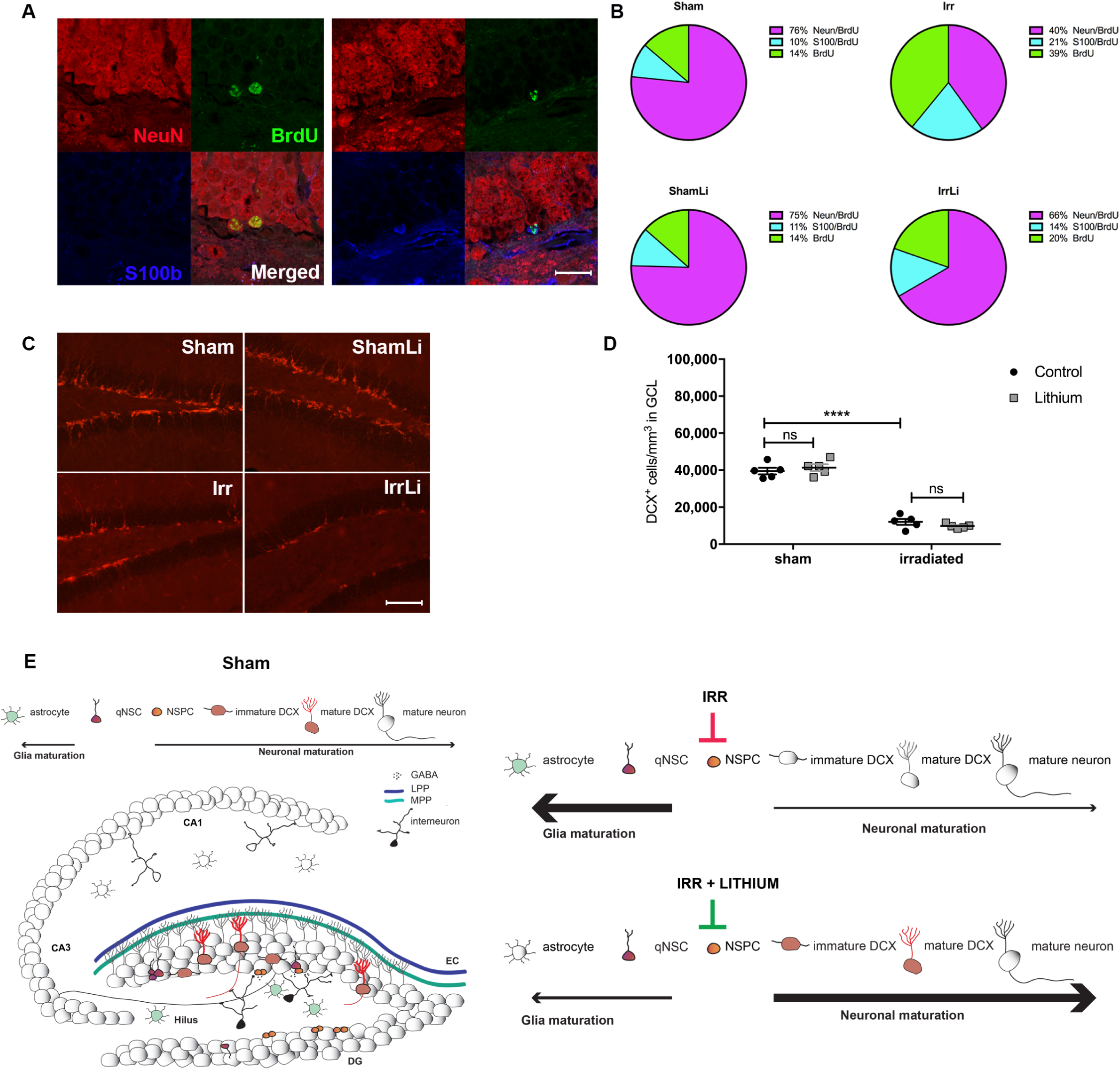
Lithium counteracts the irradiation-induced changes in NSPCs fate progression in the DG at PND105. **(A)** Representative confocal images of NeuN (red), BrdU (green) and S100β (blue) immunoreactivity in the GCL at PND105 depicting colocalization of BrdU and NeuN in mature neurons (upper panel) and colocalization of BrdU and S100β in an astrocyte (lower panel). Scale bar=20 µm. 63x magnification. **(B)** Pie chart of the percentages of BrdU cells double labeled for NeuN or S100β. The percentages of NeuN+/BrdU+ cells were significantly increased by lithium in the irradiated group (\*\**p*=0.0031) but not in sham (*p*>0.9999). 2-way ANOVA: irradiation (F_1,14_=22.5, \*\*\**p*=0.0003), LiCl (F_1,14_=7.02, \**p*=0.0191). The percentages of S100β+/BrdU+ were significantly increased after irradiation but not altered by lithium treatment. 2-way ANOVA: irradiation (F_1,14_=4.89, \**p*=0.0442), LiCl (F_1,14_=0.831, *p*=0.3774). The remaining percentages of BrdU+ cells were significantly increased in the irradiated group (\**p*=0.0137) but not altered in the other groups. 2-way ANOVA: irradiation (F_1,14_=13.1, \*\**p*=0.0028), LiCl (F_1,14_=5.12, \**p*=0.0401). **(C)** DCX immunoreactivity in the GCL at PND105 in different treatment groups. Scale bar=100 µm. 20x magnification. **(D)** Interleaved dot plot graph of the quantification of DCX^+^ cells in the GCL showing the effect of irradiation (\*\*\*\**p*<0.0001) on the density of DCX positive cells in the GCL. 2-way ANOVA: irradiation (F_1,16_=372.2, \*\*\*\**p*<0.0001), LiCl (F_1,16_=0.02213, *p*=0.8836). **(E)** Schematic drawing of the hippocampal network on the left. Input signals from the entorhinal cortex (EC) are carried through two connectional routes made of the axons of the medial (light green) and lateral (blue) perforant pathways, MPP and LPP respectively. These axons establish stable synapses with the dendrites of the mature granule cells neurons (grey) and weak ones with the immature doublecortin (DCX) cells (red) in the granule cell layer (GCL). At the boundary of the GCL and the hilus is the subgranular zone (SGZ), where quiescent neural stem cells (qNSC) give rise to amplifying neural progenitors (ANP) allowing the continuous neuronal re-population of the dentate gyrus (DG). The qNSC and ANPs are multipotent stem cells, capable of giving rise to astrocytes, oligodendrocytes and neurons. The input signal from the DG is relayed to the proximal Cornus Ammonis region (CA3) through the axons of the mature granule cells that form the mossy fibre projection. The signal transduction continues to the CA1 region through the Schaffer collaterals fibers and to further cortical areas. Parvalbumin (PV) interneurons in the hilus are important in modulating, through feed-back and feed-forward inhibition, the input signals and NSC proliferation and integration through the release of the neurotransmitter gamma-aminobutyric acid (GABA). **On the right above** a schematic representation of the effects of irradiation on DG. The number of astrocytes is increased while the number of ANPs is decreased. The neuronal differentiation process is decreased in favor of an astrocytic fate progression. **On the right below** a schematic representation of the effects of lithium on the irradiated DG. Lithium acts by increasing ANPs cell number and promoting neuronal fate progression as compared to astrocytic differentiation. Error bars indicate SEMs. ***p<0.001; **p<0.01; *p<0.05.

## DISCUSSION

### Lithium is effective even when introduced long after the injury

Cranial radiation therapy is a major cause of long-term complications in pediatric patients (Gatta et al. 2014). These complications include late-occurring cognitive impairments, and a negative impact on social competence (Armstrong et al. 2010; Georg Kuhn and Blomgren 2011). It has been demonstrated in animal models that preserving or promoting neurogenesis helps attenuate the cognitive deficits observed in irradiated mice and rats (Naylor et al. 2008a; Zhou et al. 2017). Progress has been made in identifying mechanisms underlying the neuroprotective effects of lithium in rodent models of brain injury, including anti-apoptotic effects (H. Li et al. 2011) (Huo et al. 2012) (Omata et al. 2008; Q. Li et al. 2010). Regenerative effects were demonstrated when lithium was shown to reduce brain tissue loss by 39% after hypoxia-ischemia (HI) in the immature brain even when it was administered 5 days after the injury (Xie et al. 2014) in a model where neuronal cell death peaks 1-2 days after HI, but the mechanisms have not been characterized. Delayed administration of lithium towards preservation of neurogenesis following whole-brain cranial irradiation has never been shown. In this study, we demonstrate that delayed, continuous lithium treatment regulates critical aspects of neurogenesis such as proliferation, differentiation, and survival of neural progenitor cells in the dentate gyrus of the irradiated mouse hippocampus.

Following irradiation-induced hippocampal injury, the neurogenesis cascade within the GCL undergoes several long-lasting changes: an increase in NSPC apoptosis, a continuous decrease in NSPC proliferation, a decreased propensity for differentiation of those NSPCs, where gliogenesis is favored over neurogenesis, and the onset of inflammation (Fukuda et al. 2004) (Huo et al. 2012) (Blomstrand et al. 2014). Hippocampal sections, obtained on PND 77, after 4 weeks of continuous lithium treatment, from irradiated mice, revealed that lithium promoted the proliferation of NSPCs, as indicated by the increased density of BrdU^+^ cells in the GCL. In a separate study, we found that lithium increased the proliferation of mouse hippocampal-derived NSPCs in vitro, and drove them much faster through the G1 phase of the cell cycle compared to control NSPCs (Zanni et al. 2015). However, in response to pro-neurogenic stimuli, such as physical exercise, enriched environment, and antidepressants, most proliferating cells in the dentate gyrus are amplifying neural progenitors (Kronenberg et al. 2003) (Encinas, Vaahtokari, and Enikolopov 2006) (Encinas et al. 2011) (Hodge et al. 2008). Hence, it is likely that lithium increases the rate of symmetric divisions of the amplifying neural progenitor population, which would be therapeutically beneficial since those cells represent a renewable source of neuronal precursors.

Why would delayed onset of lithium treatment be beneficial? One reason is that lithium treatment could be used for all cancer survivors who have already been treated and suffer from late effects. Another reason is the possible risk of protecting or stimulating remaining tumor cells. Radiotherapy remains an important component of their treatment despite its numerous side effects (Pollack and Jakacki 2011). Some of the pro-proliferative effects of lithium are based on pathways, which have been well described (Brown and Tracy 2013) but are still not fully understood. Some effects of lithium on brain tumor cells or leukemia cells have been described. While lithium might act as an anti-tumor agent in certain types of medulloblastoma (Zhukova et al. 2014; Ronchi et al. 2010), glioblastoma (Korur et al. 2009), glioma (Cockle et al. 2015) or leukemia (“Lithium Carbonate and Tretinoin in Treating Patients With Relapsed or Refractory Acute Myeloid Leukemia - Full Text View - ClinicalTrials.gov” n.d.), its action on other tumor cell types remains uncertain. Therefore, it may be beneficial to wait until the anti-cancer therapy is finished. To show as we did, that lithium can rescue neurogenesis long after irradiation, is thus an important step before validating its use in children.

### Intermittent treatment is important

We showed for the first time *in vivo* that discontinuation of lithium in the irradiated brain could restore neuronal fate progression, thereby counteracting the irradiation-induced astrocytic differentiation of NSPCs as previously described (Dranovsky et al. 2011; Schneider et al. 2013), further suggesting a role for lithium in restoring neurogenesis after cranial radiotherapy. Continuous lithium treatment with stable serum concentrations promoted NSPC proliferation but prevented neuronal differentiation. Discontinuation allowed differentiation and integration to occur over the course of 4 weeks. Hence, we surmise that a sequential therapeutic regimen, for example one month with lithium and one month without it, would be more effective in the treatment of cognitive late effects. Another advantage would be that the side effects frequently reported in patients (Singer 1981) treated with lithium could be better tolerated. This is a novel concept that deserves to be tested in patients.

### Identification of mechanisms

Studying both rat neural stem cells in vitro and mouse hippocampal neurogenesis in vivo enabled us to identify molecular mechanisms underlying the positive effects of intermittent lithium treatment. Two indices related to the functional integration of DCX+ cells into the GCL are the orientation of integrating DCX+ cells, and the maturity of their dendritic processes (Seki et al. 2007; Plümpe et al. 2006). Type-3 neuroblasts possess an elongated cell body, flanked by processes that lie tangential to the SGZ suggestive of an early stage of maturation, while newborn neurons have a radial process to the SGZ that is indicative of functional integration into the granule cell layer (Eisch et al. 2008). By phenotyping the orientation of dendritic processes in cells double-labeled with BrdU and DCX, we found as previously reported (Chakraborti et al. 2012; Naylor et al. 2008b) that irradiation decreased radial migration, yet increased the number of parallel dendritic processes in a different subset of neurons within that population of cells. In irradiated mice treated with lithium, the percent of cells with parallel dendrites was reduced but the radial migratory pattern was unaffected. Arguably, lithium protects against irradiation damage by limiting the number of cells with parallel dendritic processes so that immature neurons are not maintained in that stage. Overall, these data suggest that lithium discontinuation is pivotal for the late critical period of newborn cell survival as well as their structural and synaptic integration. Here irradiation persistently reduced dendritic complexity (number, length and area of branches) and spine density, similar to a previous study in which these changes were attributed to the overexpression of the synaptic plasticity-regulating postsynaptic density protein (PSD-95) in immature neurons in the GCL (Parihar and Limoli 2013). PSD-95 is believed to be a key regulator of dendritic morphology and when overexpressed it adversely affects dendritic morphology and complexity (Charych et al. 2006). We extended these findings and proved that continuous lithium treatment followed by a period without lithium acts positively on dendritic maturation and complexity, which is arguably a morphometric parameter that has important functional implications in memory function and anti-stress mechanisms (Besnard et al. 2018). We hypothesized that this is attributable to key regulatory proteins involved in cytoskeletal rearrangements (Tppp) and synaptic transmission (GAD65). Tppp expression is known to increase the stability of microtubule network thereby playing a crucial role in cell differentiation (Oláh, Bertrand, and Ovádi 2017), while knockdown of GAD65 impairs maturation of newborn granule cell (Overstreet-Wadiche, Bensen, and Westbrook 2006). Strikingly, both Tppp and GAD65 were upregulated, at the mRNA and protein levels *in vitro* and *in vivo* respectively, two weeks after discontinuation of lithium (PND 91) in our model, further confirming the positive modulatory effects of lithium on the neurogenic process as well as the importance to discontinue the proliferative drive of lithium to allow integration of the newborn cells and deter cancer relapse. The increased expression of Tppp and GAD65 was accompanied by a decreased in methylation of these two regulatory genes in the irradiated group that received lithium, supporting the hypothesis that the positive effects of lithium after irradiation are likely to be attributed to an epigenetic regulation of genes important to cell fate and maturation.

More importantly, these data support the novelty of the present study, showing that intermittent lithium treatment has unprecedented targets in the irradiated brain. Lithium has the potential to become the first pharmacological treatment of cognitive late effects in childhood cancer survivors.

## MATERIALS AND METHODS

### Animals and Ethical Permissions

C57BL/6 mice were obtained from Charles River Laboratories (Sulzfeld, Germany). The mice were kept under standard daylight conditions (12-hours light cycle) and provided with food and water *ad libitum*. All experiments were approved by the Animal Research Ethics Committee (Northern Stockholm committee of the Swedish Agricultural Agency) in accordance with the national animal welfare legislation. The ethical identification numbers were: N9-12 and N248-13 for the *in vivo* study, and N284/11 and N190114 for the *in vitro* study. Female mice were used for the experiments. Animals were delivered with their respective dams and were separated at weaning age of the pups (PND21).

### Irradiation Procedure

Mice were anaesthetized using isoflurane at 4 % for induction followed by 1.5–2 % throughout the procedure. Mice were placed on a custom-made Styrofoam frame in prone position (head to gantry), and the frame placed inside an X-ray system (Precision X-RAD 320, North Branford, CT, USA) setup in-house for *in-vivo* targeted radiotherapy research with an energy of 320KV, 12.5 mA and a dose rate of 0.75 Gy/min. The whole brain was irradiated with a radiation field of 2 x 2 cm. A single dose of 4 Gy was delivered to each animal on postnatal day (PND) 21. The source-to-skin distance was approximately 50 cm. The sham-irradiated controls were anesthetized but not irradiated.

### Lithium *in vivo* administration

Female littermates (5-6 animals in each cage) were randomly assigned to lithium chow (2.4 g/kg Li_2_Co_3_, 0.24%, TD.05357 Lithium Carbonate Diet 2018, Harlan laboratories, Netherlands) or control chow diet (T.2918.CS, Harlan laboratories, Venray, The Netherlands). This regimen was determined in our previous study (Zanni et al. 2017) and was sufficient to yield a lithium serum concentration of 0.7-0.9 mM in mice (Zanni et al. 2017), which is equivalent to the commonly used 0.6-1.2 mM therapeutic range in humans. The lithium chow was maintained four weeks, from PND 49 to PND 77 (**Figure 1A** *in vivo* study design).

### Immunohistochemistry

Animals were injected intraperitoneally with 50 mg/kg BrdU (5-Bromo-2’-Deoxyuridine) (B5002, Sigma-Aldrich, St. Louis, MO, USA) for five days starting from PND 72. Animals were sacrificed for analysis at three different time points, each separated two weeks apart: at PND 77, PND 91 and PND 105.

Immunohistochemistry and quantification were performed as previously described (Zanni et al. 2017). The following primary antibodies were used: rat anti-BrdU (1:500, AbD Serotec, Kidlington, UK) and goat anti-DCX (1:100, Santa Cruz Biotechnology Inc., CA, USA). The following secondary antibodies were used: biotinylated donkey anti-goat IgG (H+L) (Molecular Probes, Paisley, UK) and biotinylated donkey anti-rat IgG (H+L) (Jackson ImmunoResearch Europe, Suffolk, UK). For immunofluorescence, the following primary antibodies were used: rat monoclonal anti-BrdU (1:500, AbD Serotec, Kidlington, UK), goat anti-DCX (1:200, Santa Cruz Biotechnology Inc., Dallas, TX, USA), mouse monoclonal anti-NeuN (neuronal nuclei) (1:200, Merck Millipore, Billerica, EMD Millipore Corporation, Temecula, CA, USA) and rabbit polyclonal anti-S100β calcium binding protein Abcam, Cambridge, UK). The following secondary antibodies were used: Alexa Fluor^®^ 555 donkey anti-mouse IgG (H+L), Alexa Fluor^®^ 488 donkey anti-rat IgG (H+L) (Invitrogen, Life technologies, Carlsbad, CA, USA), Alexa Fluor^®^ 555 donkey anti-goat IgG (H+L) (Biotium, Hayward, CA, USA), and Alexa Fluor^®^ 633 donkey anti-rabbit IgG (H+L) (Biotium, Hayward, CA, USA). Sections were mounted in ProLong^®^ Gold Antifade Reagent with DAPI (#8961, Cell Signaling Technology, Danvers, MA, USA). For DCX^+^ cell analysis and density measurements, a fluorescent microscope was used and the total number of cells and contour areas were estimated using unbiased counting software (Stereo Investigator, MicroBrightField Inc.; Colchester, VT, USA). The other fluorescent analyses were conducted using a confocal microscope as previously reported (van Praag, Kempermann, and Gage 1999) (Axio Observer-Z1 with ZEN lite software, Carl Zeiss AG, Oberkochen, Germany). Only cells with an entire, clearly visible cell body were counted.

### Dendrite Reconstruction and Morphometric Analyses

Imaging of dendritic arbors at PND 77 and PND 91 was performed using a confocal microscope by acquiring images with 1 µm intervals using a 20X objective lens (X20 / 0.8 Plan-Apochromat lens, Carl Zeiss) (1024 x 1024 pixels). The entire dendrites and cell bodies of each neuron in the captured images were traced manually using Neurolucida^®^ software (MBF Bioscience, Williston, VT, USA). 3-dimensional analysis of the reconstructed neurons was performed and the total dendritic length, the dendritic complexity and the cell body area were measured using Neurolucida Explorer^®^ software (MBF Bioscience). A branch order was assigned to each dendrite and then the dendritic complexity was calculated as follows; dendritic complexity = [Sum of the terminal orders + Number of terminals] * [Total dendritic length / Number of primary dendrites] (Pillai et al. 2012). To measure the extent of dendritic arborization away from the soma at different distances, Sholl analysis (Sholl 1953) was conducted by counting the number of dendritic intersections for a series of concentric spheres at 10 µm intervals. The center of a concentric sphere was placed at the centroid of the soma. Immature neurons which satisfied the following criteria were selected for analysis; (i) fully labeled DCX+ cells at a postmitotic stage (Plümpe et al. 2006), (ii) neurons were relatively isolated from neighboring DCX-positive neurons to avoid interfering with analysis, (iii) the soma located in the subgranular zone or within the inner one-third of granular cell layer of the upper blade of dentate gyrus, to compare the neurons which are at the same stage of immaturity. Truncated cells were excluded from the analysis. To assure impartiality morphometric analysis was performed blindly by the same investigator.

### Protein quantification

Hippocampal tissue was processed for protein quantification as previously described (Osman et al. 2016). The Wes capillary electrophoresis system (Protein Simple-Bio Techne, San Jose, CA) was used for all protein quantitation. Sample aliquots were thawed and diluted to 0.2 μg/μl for all targets using 0.1x Sample Buffer and 5x Master Mix (1:1 mix of 400 mM DTT and 10x Sample Buffer) according to manufacturer’s instructions. Samples were denatured at 95 °C for 5 min. Rabbit □-GAD65 (PA5-22260, Thermo Fisher Scientific, USA) and Rabbit □-Tppp (ab92305, Abcam, USA) were purchased separately and used at a concentration of 1:200 and 1:2,000 respectively. Rabbit □-Vinculin was used as housekeeping control at a concentration of 1:200,000. An □-rabbit secondary antibody was provided in the kit and was used according to manufacturer’s instructions. For protein quantification the total chemiluminescent peak area was normalized to the respective reference capillary of the housekeeping control. The normalized peak area under the curve (AUC) was used for protein quantification.

### Embryonic cortical NSPC culture and LiCl exposure procedures

Primary cultures of NSPCs were established as previously described (Tamm et al. 2006; Ilkhanizadeh, Teixeira, and Hermanson 2007). Cells were obtained from embryonic cortices (*n*=10-12/cell preparation) dissected in HBSS (Life Technologies, Carlsbad, CA, USA) from timed pregnant Sprague Dawley rats (Harlan Laboratories, Harlan, The Netherlands), (Ethical Permit: N284/11 and N190114) at E15.5 (the day of copulatory plug defined as E0). The tissue was mechanically disrupted, and meninges and larger cell clumps were allowed to sediment for 10 min. The cells were plated at a density of 40,000/cm^2^ on dishes precoated with poly-l-ornithine and fibronectin (both from Sigma-Aldrich, St. Louis, MO, USA; Stockholm, Sweden). Cells were kept in enriched N-2 medium with 10 ng/ml of basic fibroblast growth factor (R&D systems, Minneapolis, MN, USA) added every 24 h and medium changed every alternate day to keep the cells in an undifferentiated and proliferative state. Cells were passaged every 5 days by detaching through scraping in HBSS. Thereafter, the cells were gently mixed in N-2 medium and plated at 1:4 density. To investigate LiCl (Sigma Aldrich, St. Louis, USA) effects, we exposed passage 3 (P3) NSPCs from 12 h before irradiation to LiCl (3 mM), as previously described (Zanni et al. 2015). A photon ^60^Co irradiation source was used to expose the NSPCs at a set distance of 80 cm and an absorbed dose of 2.5 Gy. P3 cells were harvested 24 hours after irradiation for gene expression analysis (**Figure 3A**, *in vitro* study design).

### RNA, cDNA and RT-qPCR

For real time qPCR, total RNA from culture NSPCs was extracted using RNeasy Mini Kit (Qiagen) and stored at −80 until further use. Integrity and concentration of extracted RNA were measured using Qubit (Thermo Fisher Scientific). cDNA was synthesized from extracted RNA using High Capacity cDNA Reverse Transcription Kit (Thermo Fisher) according to the manufacturer’s protocols. Quantitative real-time PCR was performed with Platinum SYBR Green qPCR Supermix-UDG (Thermo Fisher Scientific) together with site-specific primers. Expression levels were normalized to housekeeping gene, TATA-box binding protein (TBP) levels.

### RNA sequencing

Illumina TruSeq Stranded mRNA sample preparation kit with 96 dual indexes (Illumina, CA, USA) was used to prepare RNA libraries for sequencing, respectively 4 biological samples per condition (Sham, Sham+Li, Irradiated, Irradiated+Li), for a total of 16 samples. The protocols were automated using an Agilent NGS workstation (Agilent, CA, USA) using purification steps as described previously (Borgström, Lundin, and Lundeberg 2011; Lundin et al. 2010). Quality control was checked with 2100 Bioanalyzer (Agilent) with all 16 samples having RIN values of 8 or above. Libraries were sequenced on HiSeq 2500 (HiSeq Control Software 2.2.58/RTA 1.18.64) with a 2 x 126 setup using HiSeq SBS Kit v4 chemistry to an average depth 32.3 M reads (30.4 - 34.5). The data were briefly processed as following. FastQC/0.11.5 quality check was performed on raw sequencing reads, Star/2.5.1b was used to align the reads to the reference genome Rattus norvegicus genome Rnor_6.0 and QualiMap/2.2 was used to evaluate the quality of this alignment. Reads overlapping fragments in the exon regions were counted with featureCounts (subread/1.5.1) using default parameters, i.e. fragments overlapping with more than one feature and multi-mapping reads were not counted.

Differential expression analyses were performed under R/3.3.3 using EdgeR/3.16.5 package. Low count reads were filtered by keeping reads with at least 1 read per million in at least 2 samples. Counts were normalized for the RNA composition by finding a set of scaling factors for the library sizes that minimize the log-fold changes between the samples for most genes, using a trimmed mean of M values (TMM) between each pair of samples. Design matrix was defined based on the experimental design, genes-wise glms models were fitted and likelihood ratio tests were run for the selected group comparisons.

RNA-seq data have been deposited in the ArrayExpress database at EMBL-EBI (www.ebi.ac.uk/arrayexpress) under accession number E-MTAB-7238.

### MeDIP-qPCR

MeDIP-qPCR was performed using MagMeDIP kit (Diagenode) according to the manufacturer’s instructions. Briefly, rat NSPCs cells were lysed and the DNA was extracted using phenol:chloroform:isoamyl alcohol (25:24:1) (Sigma-Aldrich), purified using Purelink Genomic DNA kits (Invitrogen, now Thermo Fisher), fragmented using Bioruptur (Diagenode) and immunoprecipitated with the antibody anti-5’-methylcytosine (Diagenode), following MagMeDIP kit settings. DNA concentration was measured using Qubit dsDNA HS Assay Kit (Thermo Fisher). Immunoprecipitated DNA was quantified using RT-qPCR, as described above, and the temperature profile used was: 95 °C for 7 min, 40 cycles of 95 °C for 15 s. and 60 °C for 1 minute, followed by 1 minute 95 °C. Tppp, Gad2 promoter primers (Qiagen) and Methylated DNA and unmethylated DNA control primers (Diagenode) were used as internal controls (Fig. S1). The efficiency of methyl DNA immunoprecipitation was expressed as a relative to the percentage of the input DNA using the following equation:

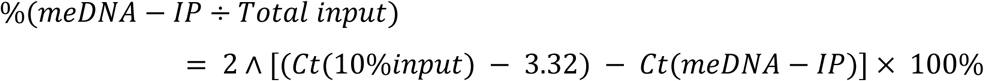

### Statistical Analysis

Statistical analysis was performed using GraphPad Prism^®^ (La Jolla, CA, USA). Statistical differences in immunohistochemistry, protein quantitation and gene expression analysis were calculated using a 2-way ANOVA analysis followed by a Bonferroni post-hoc test for multiple comparisons correction. For methylation analysis, Kruskal-Wallis test for multiple comparisons, followed by Dunn’s multiple comparisons test was performed. For dendritic analysis, linear mixed models with a random intercept for each animal, were used to account for within-individual dependencies, using R version 3.3.3 (The R Foundation for Statistical Computing, Vienna, Austria). As data were not normally distributed, natural logarithmic transformation was used to satisfy the requirement for normality. To ensure normal distributions, variables were plotted as Q-Q plots using SPSS (IBM SPSS Statistics, version 21.0, Chicago, IL, USA). For Sholl analysis, two-way ANOVA followed by post-hoc Bonferroni test was performed using GraphPad Prism (version 7.03, GraphPad Software, La Jolla, CA, USA). The data represent the mean ± SEM. In all analysis, the level of significance used was p < 0.05. All quantifications were done in a blinded fashion.

## ACKNOWLEDGEMENTS

The authors acknowledge support from the National Genomics Infrastructure in Stockholm funded by Science for Life Laboratory, the Knut and Alice Wallenberg Foundation and the Swedish Research Council, and SNIC/Uppsala Multidisciplinary Center for Advanced Computational Science for assistance with massive parallel sequencing and access to the UPPMAX computational infrastructure. This work was funded by the Swedish Research Council grant (VR-MH 2015-02675), the Children Cancer foundation grant BCF (PR2016-0129) and the Cancer foundation grant (CF CAN2015/797), grants provided by the Stockholm County Council (ALF projects), the Frimurare Barnhus Foundation in Stockholm, the Märta and Gunnar V. Philipson Foundation, the Swedish Radiation Safety Authority, the Brain Foundation, Stiftelsen Samariten, Sällskapet Barnavård. EMBO long term Fellowship (ALF 696-2013) and SSMF postdoctoral Fellowship (2015). The authors thank Ida Hed Myrberg for statistical analysis.

