## Supplementary Figure S1 for "Lithium treatment reverses irradiation-induced changes in rodent neural progenitors"

**This PDF file includes:**

Supplementary Fig. S1 and figure legend

**Fig. S1.**

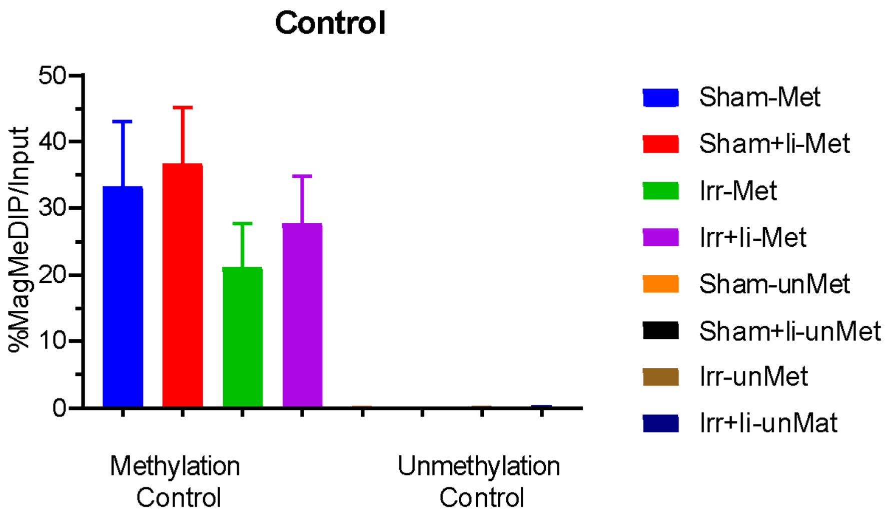


**DNA immunoprecipitation efficiency.** The efficiency of methyl DNA immunoprecipitation of the internal DNA controls of rat NSPCs included in the MeDIP assay (n = 4 - 5). For methylated control (Met), more than 20% of methylation / Input were detected for all groups (Left 4 groups). For unmethylated control (unMet), no methylated signals / Input were obtained for all groups (Right 4 groups).

**Table S1. Type or paste caption here.**

<insert Table S1 here followed by a page break>

**Additional data table S1 (separate file)**

Type or paste caption here.

### References

1.
